# High-speed, multi-Z confocal microscopy for voltage imaging in densely labeled neuronal populations

**DOI:** 10.1101/2021.12.10.472140

**Authors:** Timothy D. Weber, Maria V. Moya, Jerome Mertz, Michael N. Economo

## Abstract

Genetically encoded voltage indicators (GEVIs) hold great promise for monitoring neuronal population activity, but GEVI imaging in dense neuronal populations remains difficult due to a lack of contrast and/or speed. To address this challenge, we developed a novel confocal microscope that allows simultaneous multiplane imaging with high-contrast at near-kHz rates. This approach enables high signal-to-noise ratio voltage imaging in densely labeled populations and minimizes optical crosstalk during concurrent optogenetic photostimulation.

---

Optical measurement of activity across neuronal populations with genetically encoded calcium indicators (GECIs), such as the GCaMPs^1^, has been invaluable for studying neural dynamics with cell type specificity^2,3^. However, GECI fluorescence is nonlinearly related to neural activity, does not report subthreshold voltage changes, and has temporal resolution on the order of hundreds of milliseconds – too slow to track many of the neural processes associated with naturalistic behavior^1,4,5^. Genetically encoded voltage indicators^6-9^ (GEVIs) have the potential to address each of these shortcomings^10^. However, GEVI imaging poses two key challenges for microscope design. First, action potentials and other changes in membrane voltage occur on a time scale of milliseconds, requiring near-kilohertz sampling rates. Second, the fractional change in fluorescence of best-in-class GEVIs in response to physiological voltage fluctuations remains small – typically less than 10% – compared to widely used GECIs. Effective voltage imaging thus requires an imaging system that can provide high signal-to-noise ratio (SNR) measurements at near-kilohertz frame rates.

To meet this requirement, voltage imaging can be conducted using single-photon widefield imaging with fast (0.1–1 kHz) sCMOS cameras. However, fluorescent signal originating from the focal plane of a widefield microscope can be overwhelmed by out-of-focus background in the intact brain. To reduce out-of-focus background fluorescence, GEVIs are typically expressed in a small subset of neurons, and imaging can be performed with the aid of targeted illumination^6,7,11^. This permits widefield imaging of single neurons, although out-of-focus background still degrades the SNR of fluorescence measurements. Alternatively, high frame-rate two-photon imaging reduces out-of-focus background fluorescence but requires sparse labeling and/or is technically complex, requiring excitation powers that risk tissue heating^9,12,13^. Two-photon imaging is also incompatible with widely used rhodopsin-based GEVIs^6,8,14^.

Here, we describe a fast, volumetric confocal microscope designed for GEVI imaging and demonstrate that confocal microscopy provides several fundamental advantages over widefield microscopy. Our approach, Multi-Z Imaging with Confocal detection (MuZIC), provides both optical sectioning (background rejection) for increased SNR and simultaneous high-contrast contiguous multiplane imaging that multiplies the number of neurons that can be imaged without requiring an increase in scanning speed or laser power compared to single-plane imaging^15,16^. MuZIC permits dense three-dimensional imaging while providing resiliency to sample motion, a major concern in random-access imaging approaches^9^. We demonstrate high-SNR volumetric GEVI imaging of densely labeled neurons in mouse brain with MuZIC, both *in vitro* and *in vivo*, which has not been possible previously. Moreover, the background light rejection provided by pinholed detectors allows simultaneous voltage imaging and optogenetic stimulation with minimal optical crosstalk, thus enabling simultaneous observation and manipulation of neural activity in densely labeled populations.

In MuZIC, multiplane imaging is achieved by combining extended illumination provided by low-NA excitation with axially resolved detection obtained by high-NA fluorescence collection and detection through multiple axially distributed pinholes. Pinholes fabricated from a reflective substrate guide light emitted from outside of the focal plane of one detector to the next pinholed detector, and so forth, enabling simultaneous fluorescence detection from multiple focal planes (**Fig. 1a,b**, **EDFig. 1**). To achieve near-kilohertz frames rates for GEVI imaging, we employ a polygon scanner that provides a line rate of 117 kHz, an order of magnitude faster than resonant galvanometers. We selected pinhole diameters and spacings that minimize ‘dead space’ between planes, yielding efficient light collection throughout a 150×150×36 μm volume (12 μm inter-plane spacing) that can be imaged at a frame rate of 916 Hz while maintaining micron-scale resolution (FWHM = 2.2 μm; **EDFig. 2**). We first tested MuZIC in fixed samples containing a membrane-bound fluorescent protein. MuZIC essentially eliminates background fluorescence that otherwise degrades SNR during widefield imaging, producing high-contrast images (**Fig. 1c**).

**Figure 1 –.**
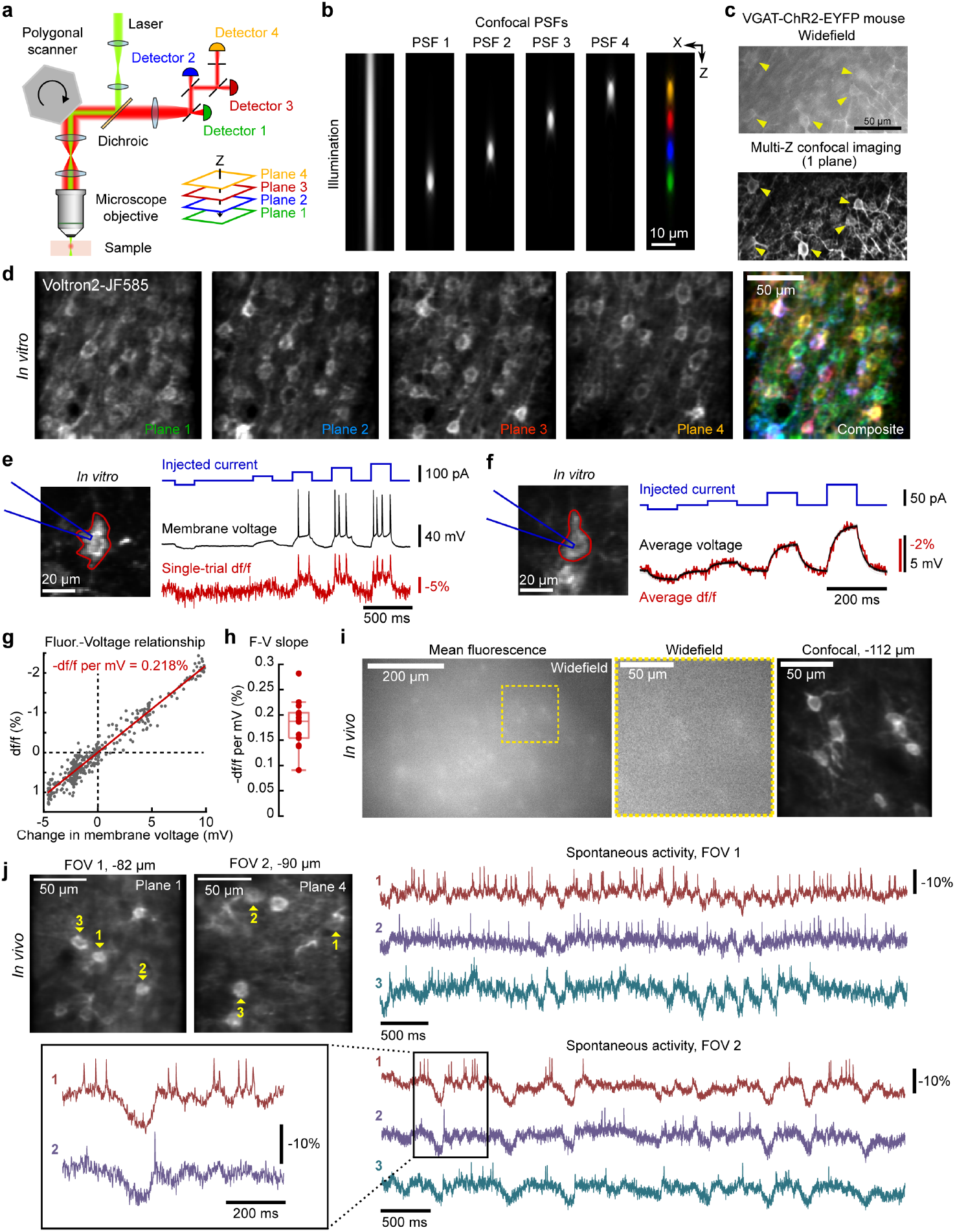
MuZIC enables high-SNR GEVI imaging in densely labeled tissue. **a.** Schematic of optical system. High speed scanning is achieved with a 128-facet polygonal scanner operating at 54,945 RPM. Excitation light (*green*) underfills the objective creating an axially extended illumination beam. Fluorescence is collected through a series of axially distributed reflective pinholes which map to different planes in the sample. **b.** Numerical simulation of illumination beam (left) and point spread functions (PSFs) associated with each image plane (*colors*). **c.** Membrane-bound YFP in a VGAT-ChR2-YFP mouse visualized with MuZIC and widefield microscopy to demonstrate optical sectioning. Confocal pinholes reject background fluorescence while preserving in-plane signal. **d.** Neurons densely labeled with Voltron2_JF585_ can be simultaneously visualized across four focal planes. **e.** Simultaneous whole-cell patch-clamp recording and voltage imaging in a cortical pyramidal neuron *in vitro*. Subthreshold membrane potential dynamics and action potentials can be detected optically in single-trial measurements. **f.** Averaging across trials reduces shot noise to reveal millivolt-scale dynamics, which are reported in a linear fashion (**g**). **h**. The relationship between changes in fluorescence and membrane potential was similar across recorded neurons (n=14 cells, 3 animals; box plot displays median and interquartile intervals). **i.** Representative widefield (*top*) and MuZIC image of excitatory neurons densely labeled with Voltron2_JF552_ (*bottom right*) acquired *in vivo*. Individual neurons are difficult to identify with widefield imaging but easily discernable with MuZIC imaging. **j**. *In vivo* imaging of subthreshold membrane potential changes and spiking in layer 2/3 pyramidal neurons reveals correlated dynamics. Two example fields-of-view are shown. Note that action potentials ride on top of subthreshold depolarizations, as expected. FOV coordinates represent axial distance from the pial surface.

Although confocal microscopy requires a similar average illumination light intensity to widefield microscopy, the requirement of point scanning in confocal microscopy necessitates much higher instantaneous illumination intensities, by a factor *A_FOV_*/*A*_0_ (~7000× in our system), where *A_FOV_* and *A*_0_ are the FOV and excitation focal spot areas respectively. This requirement could saturate the GEVI electronic excitation rate during high-speed imaging, potentially limiting the GEVI signal. For the GEVI Voltron2^14^, we estimated that the required illumination intensity for MuZIC imaging is well below the saturation level by at least an order of magnitude (**Supplementary Note 1**), and that MuZIC would therefore be readily compatible with GEVI imaging. Notably, MuZIC provides multiplane imaging ‘for free’ – there is no requirement for increased illumination intensity - and intentionally uses a focal spot diameter somewhat larger than in standard confocal microscopy further reducing the risk of fluorescence saturation.

To establish the effectiveness of MuZIC for GEVI imaging, we first labeled neurons in the mouse motor cortex with Voltron2_JF585_, and performed simultaneous whole-cell electrophysiology and MuZIC imaging *in vitro*. In densely labeled samples, thirty to forty neurons were clearly discerned across four focal planes with high contrast (**Fig. 1d**). Action potentials and subthreshold voltage fluctuations (**Fig. 1e**) induced by intracellular current injection were readily detected in single trials using the same average illumination power typically employed in widefield imaging experiments. Averaging across trials revealed small, subthreshold changes in membrane potential (**Fig. 1f**). We found a linear relationship between voltage and fluorescence in Voltron2_JF585_-labeled neurons that was consistent across neurons (**Fig. 1g,h**; 0.19 ± 0.04 % per mV, n = 14 cells).

We next sought to understand the relationship between SNR and labeling density in confocal and widefield microscopy. Confocal microscopy provides a distinct SNR advantage over widefield microscopy because it rejects background fluorescence. However, confocal microscopy requires fast photodetectors that typically provide only modest quantum efficiency (QE) and, depending on the pinhole size, partially rejects in-focus signal. Our MuZIC system employs silicon photomultiplier (SiPM) detectors (QE: ~40%) with approximately half the sensitivity of sCMOS cameras (QE: ~80%). Approximately ~1/3 of emitted fluorescence is also rejected by the pinholes. Taking these factors into account, we find that the SNR of confocal microscopy begins to surpass that of widefield microscopy when then ratio of out-of-focus background fluorescence to in-focus signal (background-to-signal ratio; BSR) is approximately 2 (**Supplementary Note 2**; **EDFig. 3**). With higher QE detectors, SNR in confocal microscopy would exceed that achievable with widefield microscopy at an even lower background level. We examined BSR in mice with Voltron2-labeled neurons *in vivo*. When only neurons in cortical layer 1 were labeled with the GEVI, a strategy that minimizes BSR by ensuring a low density of labelled cells with positions restricted to near the focal plane, we found BSR to be 1.3 ± 0.4 (mean ± s.d.; n = 18 cells), near the break-even point. In contrast, when neurons were more densely labeled, we determined BSR to be 17.7 ± 11.6 (n = 20 cells) amongst the most brightly labeled neurons (a lower bound). Together, these measurements suggest that confocal microscopy should yield SNR comparable to widefield microscopy in sparsely labeled samples while providing several-fold enhancement in SNR (**EDFig. 3**) in densely labeled samples, even when accounting for lower detection efficiency.

We tested MuZIC *in vivo* in the motor cortex of mice in which neurons were densely labeled with Voltron2 and JF552, which crosses the blood brain barrier more effectively than JF585. Following dense labeling, individual cells were difficult to resolve with widefield imaging (**Fig. 1i**, *left and middle*). In contrast, MuZIC revealed individual neurons with high contrast (**Fig. 1i**, *right*). *In vivo* imaging of layer 2 cells in awake mice revealed fluorescence transients corresponding to spontaneous action potentials in multiple neurons simultaneously as well as correlated subthreshold voltage dynamics (**Fig. 1j**). Imaging through a cranial window induces modest spherical aberration that reduces the optical sectioning strength compared to *in vitro* imaging. However, numerical simulations indicate that overall collection efficiency – and thus SNR – remains minimally affected under these conditions (**EDFig. 4**).

Finally, we evaluated the suitability of MuZIC for all-optical physiology^17,18^, in which GEVI imaging is combined with optogenetic photostimulation for simultaneous measurement and open- or closed-loop control of neural activity. While all-optical physiology represents a powerful emerging method for non-invasively interrogating neural circuit function, light used for optogenetic stimulation can corrupt voltage imaging signals if it cannot be fully separated spectrally or stimulates emission from autofluorescent species within tissue. Detection of even small amounts of stimulation light can induce significant optical artifacts on the same order as physiological changes in neuronal voltage. Unlike widefield and two-photon microscopy, which collect light indiscriminately, confocal detection rejects light emitted from sources outside of the focal area, making it well suited for all-optical physiology. The field-of-view and pinhole diameters employed in our MuZIC system yield a ~1,000-fold reduction in detection of photostimulation light compared to widefield imaging (**Supplementary Note 3**). To evaluate the potential of MuZIC for all-optical physiology, we co-expressed the optogenetic activator ChR2 and the GEVI Voltron2_JF585_ in overlapping subsets of cortical neurons and applied blue-light stimulation to simultaneously manipulate and record neural activity (**Fig. 2a**). Action potentials and subthreshold depolarizations could be readily detected across populations of neurons (**Fig. 2b-d**). Blue-light stimulation induced subthreshold depolarization and trains of action potentials in doubly infected neurons without optical artifacts (**Fig. 2b-d**). These data demonstrate the potential for all-optical physiology in densely labeled neural populations.

**Figure 2 –.**
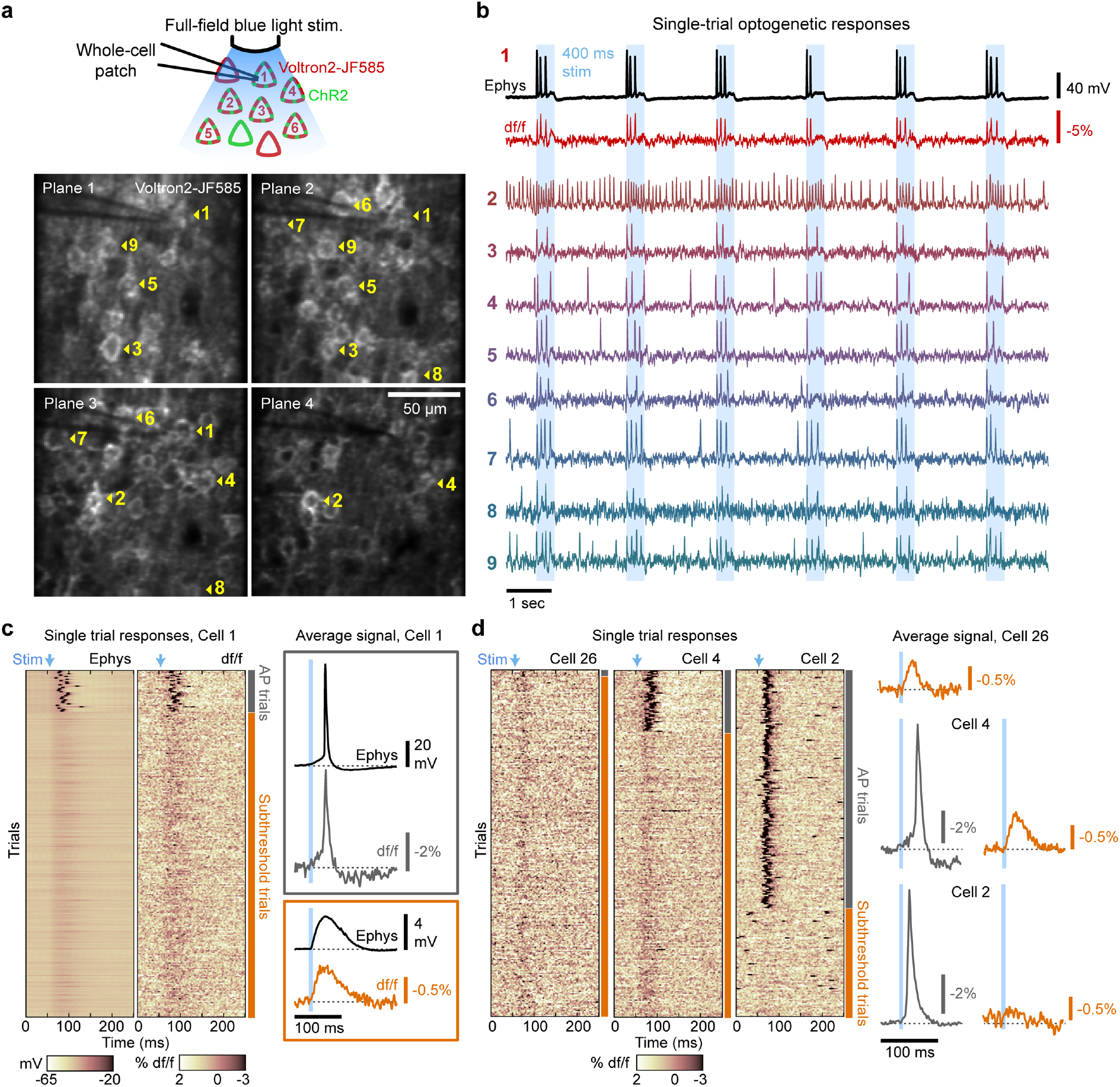
MuZIC enables simultaneous imaging and optogenetic stimulation with minimal optical crosstalk. **a.** *Top*, Schematic of experiment. Overlapping subsets of neurons are virally labeled with Voltron2_JF585_ and ChR2. Widefield optical stimulation excites ChR2-expressing neurons. *Bottom*, Expression of Voltron2_JF585_ across four imaging planes. **b**. Simultaneous electrophysiological (*black*) and optical recording (*red*) of membrane potential from cell indicated by red ROI in (***a***) as well as optical recording from eight additional Chr2+/Voltron2_JF585_+ neurons in response to repeating 400-ms blue light stimuli. **c.** Short (0.5 ms) blue light stimuli induce action potentials (top, gray bars) or subthreshold voltage deflections (*bottom, orange bars*) in both electrophysiological (*left heatmap*) and optical (*right heatmap*) recordings of membrane potential. 256 repetitions shown. Average electrophysiological and optical recordings across all trials in which action potentials (*gray box, top*) or subthreshold depolarizations (*orange box, bottom*) were elicited. **d**. Optical recordings, as in (***c***) of additional cells from the same field of view that displayed subthreshold responses (*left*) and action potentials on a minority (*middle*) and majority (*right*) of trials.

In summary, we describe MuZIC for high-contrast, volumetric GEVI imaging *in vitro* and *in vivo*. Confocal detection in MuZIC rejects background fluorescence which would otherwise prohibit widefield imaging in densely labeled samples. Fast polygonal scanners enable a near-kilohertz frame rate while extended illumination combined with an array of pinholed detectors allow multiple focal planes to be imaged simultaneously without the need for additional excitation power. These features multiply the number of neurons that can be imaged and reduce sensitivity to axial sample motion. Pinholed detectors also minimize artifacts associated with simultaneous optogenetic stimulation by three orders of magnitude, thus facilitating all-optical physiology. MuZIC systems can be built with readily available optical components for a total cost on par with commonly used widefield imaging systems built around sCMOS cameras. Additionally, we establish that confocal imaging provides a fundamental benefit in SNR over widefield imaging of all but the most sparsely labeled samples and provides a many-fold increase in SNR in densely labeled thick tissues.

## AUTHOR CONTRIBUTIONS

TDW, MVM, JM, and MNE conceived of the project. MNE and JM supervised research. TDW, JM, and MNE designed the MuZIC system. MVM and MNE prepared samples. TDW, MVM, and MNE performed experiments. TDW, MVM, and MNE analyzed data. TDW performed simulations. TDW, MVM, JM, and MNE wrote the manuscript.

## ACKNOWLEDGMENTS

The authors thank Ahmed Abdelfattah, Amrita Singh, Gabe Murphy, and Karel Svoboda for technical advice and helpful discussions. The authors thank Eric Schreiter and Ilya Kolb for sharing Voltron2 reagents pre-publication. The authors thank Michael Giacomelli for advice on silicon photomultipliers. We thank Tim Wang and David Kleinfeld for helpful comments on the manuscript. This work was supported by NIH R01EB029171.

---

**Extended Data Figure 1 –.**
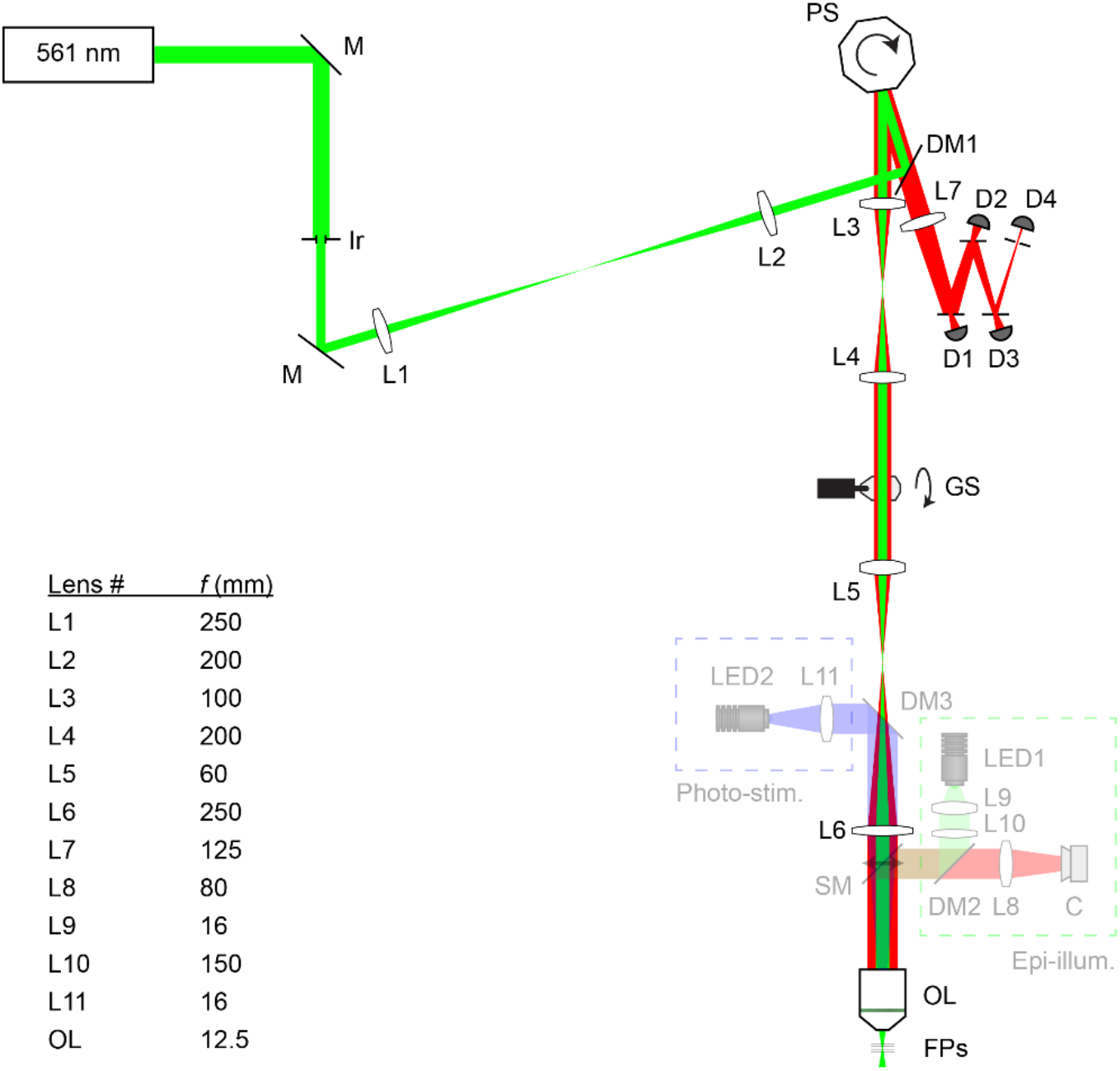
Detailed Multi-Z Confocal Microscopy (MuZIC) system schematic. The light source is a continuous-wave laser diode (“561 nm”). Blue dashed box (“Photo-stim.”) denotes components of the full-field photo-stimulation module. Green dashed box (“Epi-illum.”) denotes components of the epi-illumination widefield fluorescence microscopy unit. M: mirror, Ir: iris, L: lens, DM: dichroic mirror, PS: polygonal scanner, GS: galvanometer scanner, SM: switchable mirror, OL: objective lens, FPs: focal planes, D: detector, C: camera.

**Extended Data Figure 2 –.**
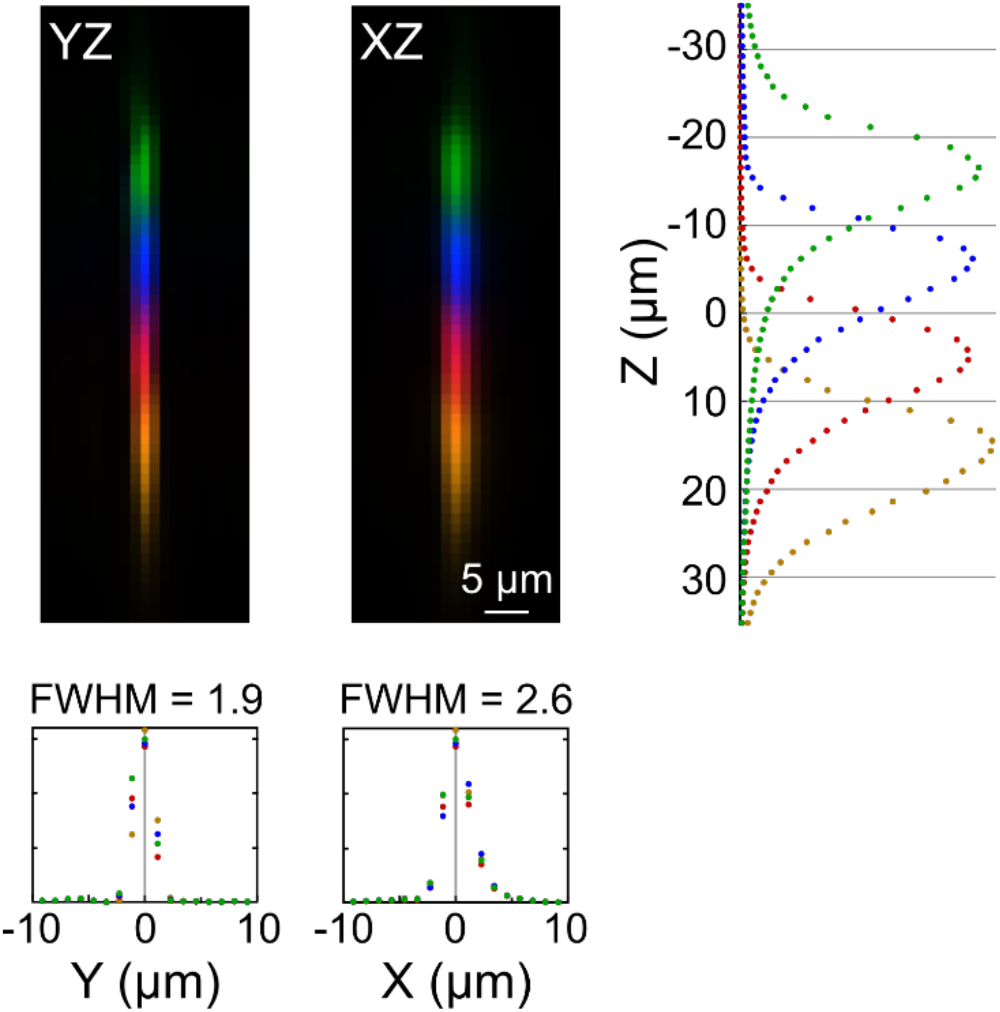
System resolution. Experimental measurements of the combined excitation and emission point spread functions (PSFs) corresponding to the four simultaneously acquired image planes. Confocal pinhole diameters were selected so that PSFs corresponding to adjacent detectors overlap, eliminating gaps between the four imaging planes, which together cover ~40 μm in the axial direction. Lateral resolution was somewhat anisotropic on account of the slower detector fall time. (X: 2.6 μm full-width half-max; Y: 1.9 μm).

**Extended Data Figure 3 –.**
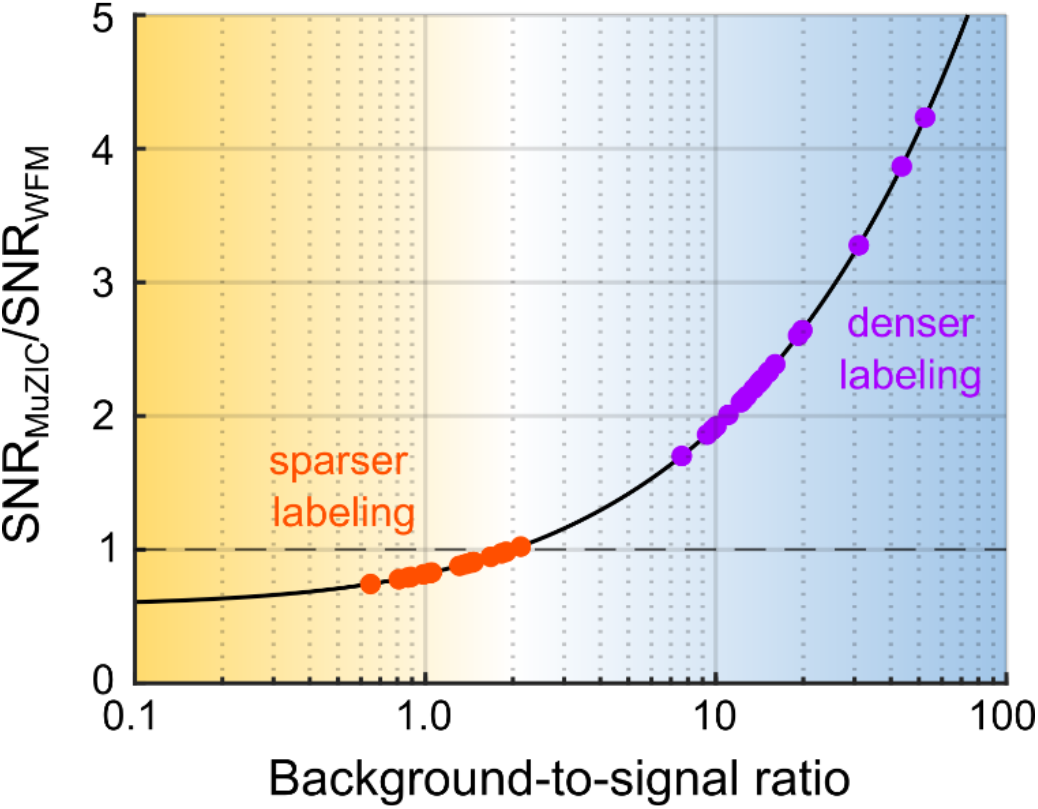
Comparison of Multi-Z Confocal Microscopy (MuZIC) and widefield fluorescence microscopy (WFM). Signal-to-noise ratio of MuZIC relative to WFM increases with increasing background-to-signal ratio (BSR). SNR of MuZIC is mostly insensitive to out-of-focus background fluorescence, while SNR of WFM decreases with increasing background. When BSR is greater than about 2, SNR of MuZIC is higher than SNR of WFM (assuming 33% detection efficiency of MuZIC relative to WFM; see **Supplementary Note 2**). The BSR associated with *in vivo* imaging of densely-labeled neuronal populations (*purple points*) was 17.7 ± 11.6 (mean ± s.d.; n = 20) for neurons with the highest Voltron2_JF552_ expression. More dimly labeled neurons with higher BSR could not be discerned from background. This analysis suggests a lower bound for SNR_MuZIC_/SNR_WFM_ of approximately 2.25 in this regime. BSR associated with sparse labeling was near unity (*orange points*; 1.3 ± 0.4; n = 18).

**Extended Data Figure 4 –.**
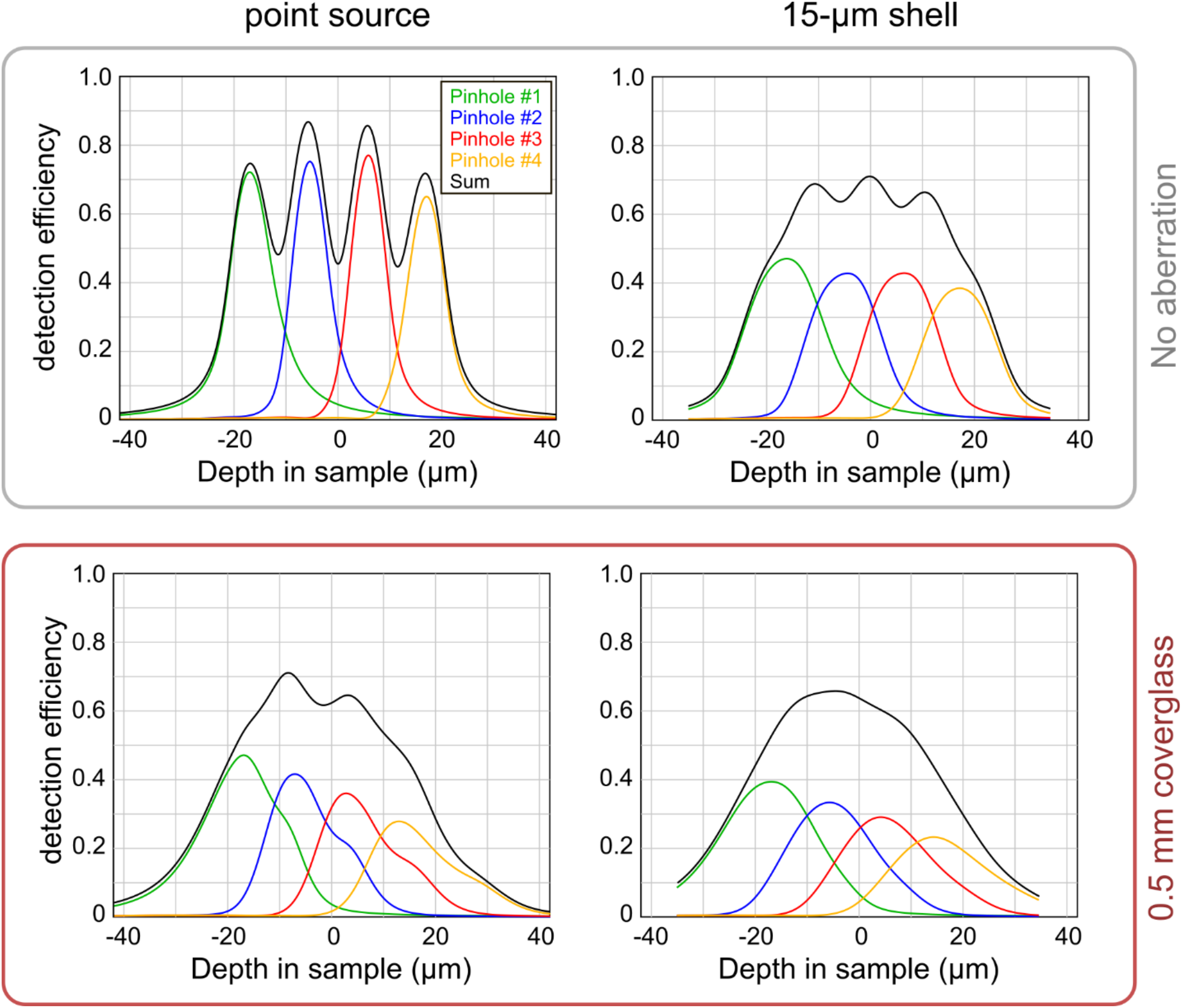
Confocal collection efficiency. Simulated confocal collection efficiency for fluorescence emitted from point sources (*left*) and 15-μm shells (*right*) located at different axial positions. Axial resolution is degraded as a result of spherical aberration induced by imaging through 0.5 mm cover glass (*bottom*; RI = 1.52). Lateral resolution is not degraded as much as axial resolution, meaning that the total fluorescence together collected by all detectors is only modestly reduced. See **Methods** and **Supplementary Note 4** for more details.

## ONLINE METHODS

### Microscope design

We designed the MZCM system (**Fig. 1a, EDFig. 1**) for imaging longer-wavelength JF585 to open the blue spectral window for optogenetic stimulation, reduce tissue autofluorescence, increase penetration depth, and match the fluorescence emission to peak detector sensitivity (600 nm). A 561 nm diode laser (Vortran Stradus 561-50) was used as an excitation source in all experiments. To underfill the objective back aperture, the laser beam diameter was reduced with an iris aperture and reimaged onto a polygonal scanner, which performed the fast axis (x) scanning. The laser beam was coupled into the optical path just before the polygonal scanner via a dichroic mirror (Chroma T570lpxr). The polygon scanner (Lincoln Laser SA-24 with DT-128-250-025-AA protected gold mirror) features 128 facets, rotating at 54,945 RPM, which generated a fast axis scan rate of 117.2 kHz. From the polygonal scanner, the beam was relayed to a galvanometer scanner (Cambridge Technology 6215H), providing the slow axis (y) scan, and relayed again to the back aperture of the objective lens. To generate an axially extended illumination focus, the effective illumination NA was limited to approximately 0.15. All experiments were performed with a 16× 0.8 NA water-immersion objective (Nikon CFI75 LWD 16X W).

Emitted fluorescence was epi-collected, de-scanned, and separated from reflected illumination light with the dichroic mirror and an additional emission filter (Chroma ET570lp). The de-scanned fluorescence was focused onto a set of axially distributed reflective pinholes (Edmund Optics #45-604, laser ablated by National Aperture). Based on the total estimated lateral magnification (83×) the projected image of each pinhole was 3.6 μm in diameter. The pinholes thus captured the majority of the fluorescence generated from the 4 focal planes. The distance between the focal planes in the sample is given by *z_p_n/M*^2^ where *z_p_* is the axial distance between the pinholes in image space, *n* is the refractive index of the sample immersion medium (1.33 for water), and *M* is the lateral magnification. In our case, *z_p_* ≈6 cm, resulting in a 12 μm interplane spacing in the sample. Care was taken to minimize the angle of incidence (< 17°) on the pinholes such that the projection of the pinhole did not appear elliptical to the oncoming beam.

Red-shifted silicon multipliers (SiPMs, Hamamatsu S14420-1550MG) were selected for their high effective photon detection efficiency (~40%). Each detector was soldered to a custom PCB which included a transimpedance amplification circuit (https://github.com/tweber225/simple-sipm). A reverse bias voltage of 50V was provided with a programmable DC power supply. Amplified signals were digitized and binned to 15 MHz (NI-5734 and NI PXIe-7961) to achieve square, approximately Nyquist-sampled pixels (1.1 μm pixel pitch vs. 2.2 μm PSF mean FWHM).

The system was also equipped with an epi-illumination widefield fluorescence microscopy module for sample navigation. The widefield path could be made available by sliding a repositionable mirror in the scanning path (Thorlabs OPX2400). The module included a green LED (Thorlabs M565L3), a similar filter set (Chroma set 49017), and a CMOS camera (Thorlabs CS165MU), and provided an approximately 0.8×0.6 mm FOV. For *in vitro* experiments, this camera was also combined with an oblique transillumination condenser and a NIR LED to provide phase-gradient contrast for whole-cell patch clamp recordings. Full-field optogenetic stimulation was provided with a blue LED (Thorlabs M470L4) coupled into the scan path via an excitation filter and dichroic mirror (Thorlabs MF475-35 and DMLP505R).

### Microscope operation

Scanning and confocal image generation were controlled with ScanImage (Pologruto et al., 2003) in combination with Wavesurfer (https://wavesurfer.janelia.org/) to synchronize current injection and optogenetic stimulation. The fast axis scan range was 150 μm (nearly 100% duty cycle) divided into 127 pixels. The slow axis (linear galvanometer) was run in a vertical bi-directional mode. Odd frames were acquired by scanning lines top to bottom, and even frames acquired by scanning bottom to top, a strategy adopted to minimize vertical flyback deadtime. The slow axis scan range and number of lines was matched to the fast axis, yielding 150×150 μm 4-plane imaging at 916 Hz. One additional line scan is required for internal ScanImage synchronization, which brought the total number of fast scans per frame period to 128. Matching the fast scans per frame period to a multiple of the polygon facet count eliminates frame-to-frame variation in each acquired line that results from small variations in facet-to-facet pointing and polishing.

For *in* vitro experiments, imaging laser (561 nm) power was set to 0.2 - 0.9 mW. Over the 0.0225 mm^2^ FOV, this is an irradiance 9 – 40 mW/mm^2^, well below the molecular saturation limit for JF585 (**Supplementary Note 1**). More power (2 – 6 mW) was required for *in vivo* experiments most likely due to reduced transparency of the optical window and increased absorption by blood and scattering due to deeper imaging. Full-field optogenetic stimulation intensity was adjusted with a neutral density filter and/or LED current control.

### Animals

All animal procedures and experiments were carried out with approval from the Boston University Institutional Animal Care and Use Committee (IACUC) and in accordance with National Institutes of Health (NIH) policies and guidelines. Both male and female C57Bl/6J mice (JAX #000664) and Emx1-IRES-Cre mice (JAX #005628) were used in this study. **Figure 1c** also included fixed tissue from a VGAT-Chr2-EYFP mouse (JAX #014548).

### Adult Viral injection

Anesthesia was induced using 5% isoflurane in O_2_ and was maintained during surgical procedures using 1-2% isoflurane in O_2_. Bupivacaine (0.1 ml, 0.5%) was injected under the skin covering the skull. The skin and periosteum covering the skull were removed and the skull thinned overlying the sites of viral injection(s). For all injections, virus was injected using a manual volume displacement injector (MMO-220A, Narishige) connected to a glass pipette (5-000-2005, Drummond Scientific) pulled to a 30 μm tip (P-2000, Sutter Instruments) that was beveled to a sharp tip. Pipettes were back-filled with mineral oil and virus was front-loaded before injection. Pipettes were inserted through the thinned bone to the appropriate depth and virus injected at 20 nl min^−1^. Animals were injected at 7-9 weeks of age.

Animals were unilaterally injected with a 1:10 dilution of AAV1-hSyn-FLEX-Voltron2-ST-WPRE virus (E. Schreiter) in ALM (coordinates in mm from Bregma: AP +2.5, ML +1.5, DV −1.2 and −0.6) and M1 (from Bregma: AP + 0.75, ML +1.5, DV −1.2 and −0.6). Cortical injections to adult C57Bl/6J mice additionally included a 1:250 dilution of rAAVretro-hSyn-Cre (Addgene #105553-AAVrg) to induce expression of Voltron2. For optogenetic experiments, AAV8-Syn-ChR2(H134R)-GFP-WPRE (Addgene #58880-AAV8) was injected into motor cortical sites at a 1:1 mixture with Voltron2 virus, or into the motor thalamus VM/VL (from Bregma: AP −1.6, ML + 1.0, DV −4.25 and 3.5). Total injection volume at each target site was 100 nL. Mice were administered post-operative sub-cutaneous injections of ketoprofen (5 mg/kg) and buprenorphine (0.1 mg/kg) in saline for pain management. Viruses were allowed to express for 3-4 weeks before imaging was performed.

### Neonatal viral injection

Intraventricular viral injections were carried out in C57Bl/6N (Charles River Labs) neonates^1^. Briefly, postnatal day 1 (P1) pups were anesthetized by placing them on an aluminum plate resting on ice. A kimwipe was used to prevent direct contact of the pup with the cold plate. When the pup was no longer responsive to toe pinch, it was moved onto the stereotaxic apparatus while maintained cold. Injections were targeted to the right lateral ventricle (approximate coordinates in mm from Lambda: AP +1.4, ML +0.8, DV −1.5). 1 μL of solution containing a total of 2.0×10^9^ g.c. of Voltron2-ST virus and 1.0×10^9^ g.c. of Cre virus in sterile PBS was injected slowly into the ventricle. For optogenetic experiments, 3.0×10^9^ g.c. of Chr2(H134R)-GFP virus were also included in the injection solution. To visualize successful delivery of virus, 0.05% Trypan blue was added to the injected solution. Pups were allowed to recover on a 37°C warming pad before being returned into the cage with the dam. The behavior of the dam was carefully monitored for 2 hrs following injections to ensure maternal care of the pups.

### In vitro imaging and electrophysiology

Acute slice preparation was carried out using protocols optimized for adult tissue^2^. Briefly, animals were perfused with 15mL chilled and carbogen-bubbled (5% CO_2_/95% O_2_) NMDG aCSF (in mM: 92 NMDG, 2.5 KCl, 1.25 NaH_2_PO_4_, 30 NaHCO_3_, 20 HEPES, 25 glucose, 2 thiourea, 5 Na-ascorbate, 3 Na-pyruvate, 0.5 CaCl_2_·4H_2_O and 10 MgSO_4_·7H_2_O, pH 7.3-7.4, 300-310 mOsm) at a rate of 10 mL/min via manual cardiac perfusion. Brain tissues were quickly dissected and sliced to 300 μm using a vibratome (Precisionary Instruments) in chilled/bubbled NMDG aCSF. Following Na+ spike-in in 37°C NMDG aCSF, slices were placed in room-temperature HEPES aCSF (in mM: 92 NaCl, 2.5 KCl, 1.25 NaH_2_PO_4_, 30 NaHCO_3_, 20 HEPES, 25 glucose, 2 thiourea, 5 Na-ascorbate, 3 Na-pyruvate, 2CaCl_2_·4H_2_O and 2 MgSO_4_·7H_2_O, pH 7.3-7.4, 300-310 mOSm) containing 25 nmol of JF585^3,4^ and incubated for 1 hour in the dark. Slices were then moved to HEPES aCSF for 1 hr to washout excess dye before imaging. *In vitro* slices were imaged while perfusing bubbled, room temperature HEPES aCSF. Whole-cell recordings were made using filamented glass pipettes (Sutter #BF150-86-10) pulled to 3-8 MOhm resistance (Sutter P-1000 Micropipette Puller), and intracellular recording buffer containing (in mM) 145 K-Gluconate, 10 HEPES, 1 EGTA, 2 Mg-ATP, 0.3 Na_2_-GTP, and 2 MgCl_2_ (pH 7.3, 290-300 mOsm). A patch-clamp headstage (Molecular Devices #1-CV-7B) mounted on a motorized 4-axis Siskiyou MX7600 manipulator, and Axon Instruments MultiClamp 700b amplifier were used for all recordings.

### In vivo cranial window implant and labeling

Adult mice that received Voltron2-ST injections as neonates (see above) were used for *in vivo* imaging experiments. At 5-6 weeks, a 2.5 mm-diameter craniotomy was made above the motor cortex. Cranial windows were fabricated by stacking two 2.5mm coverglass discs (#1.5 glass, Potomac) and one 3.5mm disc (#1.5 glass, Potomac) and secured them together with optical glue. These glass windows were implanted into the craniotomy and affixed to the skull with super glue. A headbar was then affixed caudal to the window. Dental acrylic (Jet repair; Pearson Dental) was used to secure the headbar to the skull and protect exposed bone. 24 hours prior to imaging, a retroorbital injection was performed to deliver 100 μL of solution containing 20 μL of Pluronic F-127 (20% w/v), 20 μL of DMSO, and 100 nmol of JF552 dye in sterile PBS.

### Image processing and analysis

All data were analyzed using custom scripts written in MATLAB (Mathworks). Binary regions of interest (ROIs) spanning all four planes were determined semi-automatically^5,6^ using the average images measured across all even (forward scan) frames. ROIs determined from even frames were warped onto odd frames (reverse scan frames) using an affine transform computed using a similarity-based metric applied to averages of odd and even frames to correct for small differences induced by bidirectional scanning. ROIs were not constrained to be spatially contiguous. The fluorescence time series corresponding to each ROI was computed as the summed intensity over all pixels included within the ROI. In all analyses, df/f was calculated as (f-f_0_)/f_0_, where f_0_ was taken to be the mean fluorescence. The relationship between df/f measurements and voltage measurements were determined with least squares fits (**Fig. 1g,h**). In **Fig. 2d,e**, single action potentials (measured both electrically and optically) were peak-aligned in time prior to averaging. In **Fig. 1h**, repeated measurements (using subthreshold and suprathreshold current steps) were made in some cells. In some experiments, a modest amount of photobleaching (typically less than 20%) was corrected for with a high-pass 2^nd^ order Butterworth filter with a cutoff frequency of 0.5 Hz. Traces were low pass-filtered using causal boxcar filters. Lateral motion correction was applied to image stacks recorded in vivo. Displacements in the x- and y-directions were determined using cross-correlation at subpixel resolution. Displacements were calculated separately for each image plane and the mean displacement applied uniformly to all planes. In **EDFig. 3**, measurements of background-to-signal ratio for sparse in vivo labeling were calculated based on images in reference [3]. Somata locations were identified manually and fluorescence was taken as the median pixel value within a 7.5-pixel radius around those locations. Background fluorescence was taken as the median fluorescence across four same-sized ROIs displaced to the left, to the right, above, and below the soma center by 50 pixels.

### PSF simulations

To estimate the collection efficiency and effect of cranial window-induced aberrations, we developed a wave-based numerical model for the MZCM point spread function (PSF). We computed the illumination and detection 3D PSFs separately, taking into account upstream pinhole occlusion (see **Supplementary Note 4** for additional details). The illumination and detection PSFs are pair-wise multiplied to yield the final confocal PSF_s_.

## DATA AVAILABILITY

Primary and derived data described in this study will be made available on Figshare upon publication.

## CODE AVAILABILITY

All code used for data analysis and design information related to construction of our MZCM imaging system will be made available via Github.

## Supplementary Note 1 – Fluorophore saturation in confocal microscopy

We define:

*p* = illumination power
*I* = illumination intensity
*A_FOV_* = area of field of view
*A*_0_ = area of laser focus (in case of confocal microscopy)
*τ_f_* = GEVI fluorescence lifetime (or excited-state lifetime)
*σ_a_* = GEVI molecular absorption cross-section

We compare confocal and widefield microscopy, assuming that the illumination power delivered to the FOV is equal in both cases (i.e. *P_c_* = *P_W_*), and that acquisition time per FOV is the same in both cases. To achieve the same overall fluorescence signal per GEVI molecule, we must have

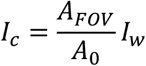

That is, the intensity of the focused laser in confocal microscopy must be significantly higher than that of the defocused illumination in widefield microscopy. As such, confocal microscopy runs the risk of saturating the GEVI molecules.

Saturation occurs when the excitation rate into the GEVI excited state exceeds the decay rate out of the excited state. That is, saturation occurs when

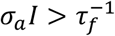

This defines a saturation intensity given by

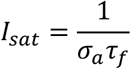

To calculate *I_sat_* for JF585, we use the following numbers:

Extinction coefficient [reference]: *ε*_0_ = 109,000 M^−1^ cm^−1^
Fluorescence lifetime [reference]: *τ_f_* = 3.8 × 10^−9^ s

From the relation^1^ 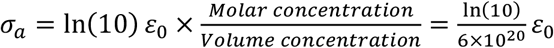. we have for JF585: *σ_a_* = 4 × 10^−16^ cm^2^ = 4 × 10^−8^ μm^2^

The saturation intensity for JF585 is thus

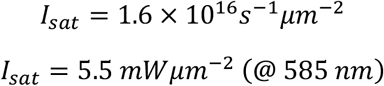

To avoid saturation with our confocal microscope, we must satisfy the condition:

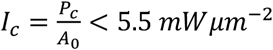

In our case, *A*_0_ ≈ 4 *μm*^2^, meaning we must satisfy the condition *P_c_* < 22 *mW*

We typically used laser powers *P_c_* at the sample in the range 0.4-0.8 mW. In other words, we were well below the saturation limit by more than an order of magnitude.

## Supplementary Note 2 – SNR in widefield and confocal microscopy

Here we compare the SNRs associated with voltage imaging for confocal and widefield microscopy, and examine how these depend on sample density.

Let *F* be the GEVI fluorescence produced in the sample (#photons/s). The measured fluorescence is then given by 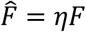, where *η* is the overall detection efficiency. We assume the fluorescence is linearly dependent on the voltage *V*. That is

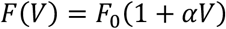

where *F*_0_ is the baseline fluorescence. The measured voltage is thus given by

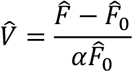

and the noise associated with the measured voltage is

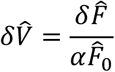

Where we have assumed that the measurement of 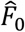 is noiseless since it can be acquired over long times.

We thus arrive at the general result

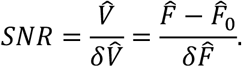

At this point, we can divide the fluorescence into in- and out-of-focus components, each detected with associated detection efficiencies. That is

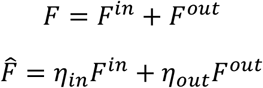

The baseline fluorescence is assumed large enough that the measurement is shot-noise limited. We further assume that the voltage variations of interest are confined to the in-focus fluorescence. We thus have

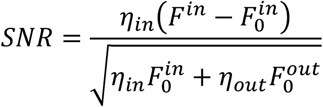

Leading to the general result

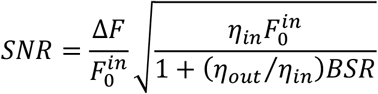

where 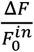 is the voltage response of the GEVI, and 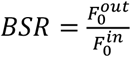 is the background to signal ratio, which characterizes the sample density (higher *BSR* corresponds to higher density).

We can now compare the *SNRs* associated with confocal versus widefield microscopy.

For a confocal microscope (*η_in_* = *η_c_*; *η_out_* = 0),we have

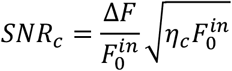

For a widefield microscope (*η_in_* = *η_out_* = *η_w_*), we have

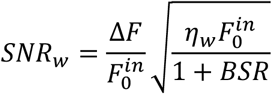

Hence

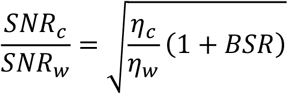

When imaging a point fluorescent object, the ratio of (in-focus) detection efficiencies between confocal and widefield microscopy is dominated by the quantum efficiencies of the associated detectors (SiPM versus sCMOS camera). That is, 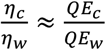, corresponding in our case to about 0.5. However, when imaging neuronal somas which resemble 15 μm shells, the ratio 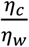 is reduced somewhat because the fluorescence generated between the focal planes is partially rejected by the pinholes (see Fig. S4). We use the estimate obtained from simulations 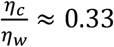, leading to the plot shown in Fig. S3.

## Supplementary Note 3 – Rejection of optogenetic stimulation light

We define:

*O* = background generated during optogenetic stimulation (either by leakage of optogenetic excitation into fluorescence detection channel, or by crosstalk generation of GEVI fluorescence)
*A_FOV_* = area of field of view
*A_p_* = area of pinhole projected into sample (in case of scanned, pinholed detection)
*η_w_, η_c_* = overall detection efficiencies for widefield and confocal microscopies respectively

In the case of widefield imaging, the detected background is given by

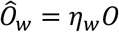

In the case of scanned detection through a pinhole, the pinhole by itself provides no optical sectioning, meaning that in- and out-of-focus background are equally detected as in widefield imaging. However, it does reduce the amount of time over which the background is collected by the ratio *A_p_* / *A_FOV_* (assuming that the optogenetic stimulation duration is longer than a frame time). Note that this same ratio applies for all depths since in- and out-of-focus background are detected with equal efficiency per depth slice.

Hence, the optogenetic-induced detected background in the case of confocal GEVI imaging is given by

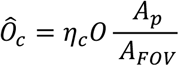

In our case *A_FOV_* = 22500 μm^2^ and *A_p_* 10 μm^2^.

Also 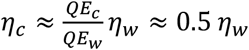.

Hence

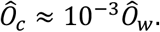

## Supplementary Note 4 – Simulation details

The 3D PSFs were calculated using mixed spatial frequency domain coordinates—that is spatial frequency coordinates (*κ_x_*, *κ_y_*) for transverse dimensions and a real space coordinate (*z*) for the axial dimension. Computation in all real space coordinates (*x, y, z*) is also valid, but the mixed domain method has the advantage that defocus and pupil aberrations can be represented easily in the transverse spatial frequency domain.

To calculate the illumination PSF, we generated a discretized binary pupil with radius *κ*_max_ = NA_*i*_/λ*_i_* (NA_*i*_ = 0.15, λ*_i_* = 0.561 μm). The pupil was zero-padded to yield a pixel spacing of 0.167 μm. This discretized function represents the Fourier transform of the illumination field, also called the radiant field, in the sample focal plan. To extend to 3D sample space, the in-focus radiant field is replicated along the 3^rd^ dimension (*z*). Each slice of the stack is multiplied by the frequency domain representation of defocus, 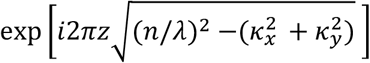, where z ranged from −64 to +64 μm with 0.167 μm spacing, and *n* is the sample index of refraction. Each slice is 2D discrete Fourier transformed and multiplied with its complex conjugate, yielding the 3D illumination PSF (**Fig. 1b**, left).

Calculation of detection PSFs is more complicated due to upstream pinhole occlusion. Our general strategy was to compute the 3D PSF for each discretized point making up the detection pinhole (pinhole points) and later sum each to form a final detection PSF. This simulates the pinhole’s admittance of intensity (incoherent summation). Pinhole points must be calculated separately because they experience different degrees of occlusion depending on their transverse position.

For each pinhole point, we initialize an array with the same size as the illumination pupil function. We set the point’s value to 1 with all other locations equal to 0. This array is 2D Fourier transformed and becomes the radiant field representation of the pinhole point. For points in pinholes subject to occlusion (channels 2-4), we numerically propagated the radiant field to the upstream pinhole’s axial location using the aforementioned defocus function. To simulate an upstream blocking pinhole, we invoked the convolution theorem and convolved the propagated radiant field with the Fourier transform of the inverse of the pinhole. We then repeated this process for any additional upstream pinholes and finally propagated the field back to the system focal plane. To simulate imaging, we multiplied the field at the focal plane by a discretized binary pupil of radius *κ*_max_ = NA_*d*_/*λ_d_* (NA_*d*_ = 0.8, *λ_d_* = 0.61 μm). We then followed the same approach used for illumination to extend to 3D and compute the 3D PSF. This entire process repeated for all points comprising the pinhole and finally summed to yield each channel’s 3D detection PSF.

Cranial window aberrations are introduced by adding the following phase profile to the circular pupil for both illumination and detection PSFs,

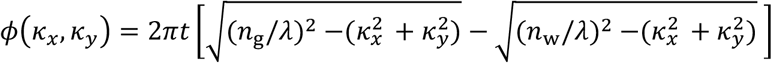

In the expression, *t* is the thickness of the glass and *n*_g_/*n*_w_ are the refractive indices of glass and water. When extending either PSF to 3D, the defocus function’s z range is adjusted to account for the shift in focal plane induced by the refractive index mismatch.

## REFERENCES

1. Chen, T.-W. et al. Ultrasensitive fluorescent proteins for imaging neuronal activity. Nature 499, 295–300 (2013).

2. Grienberger, C. & Konnerth, A. Imaging calcium in neurons. Neuron 73, 862–885 (2012).

3. Luo, L., Callaway, E. M. & Svoboda, K. Genetic Dissection of Neural Circuits: A Decade of Progress. Neuron 98, 256–281 (2018).

4. Wachowiak, M. et al. Optical dissection of odor information processing in vivo using GCaMPs expressed in specified cell types of the olfactory bulb. J. Neurosci. 33, 5285–5300 (2013).

5. Wei, Z. et al. A comparison of neuronal population dynamics measured with calcium imaging and electrophysiology. PLoS Comput Biol 16, e1008198 (2020).

6. Abdelfattah, A. S. et al. Bright and photostable chemigenetic indicators for extended in vivo voltage imaging. Science 365, 699–704 (2019).

7. Adam, Y. et al. Voltage imaging and optogenetics reveal behaviour-dependent changes in hippocampal dynamics. Nature 569, 413–417 (2019).

8. Piatkevich, K. D. et al. Population imaging of neural activity in awake behaving mice. Nature 574, 413–417 (2019).

9. Villette, V. et al. Ultrafast Two-Photon Imaging of a High-Gain Voltage Indicator in Awake Behaving Mice. Cell 179, 1590–1608.e23 (2019).

10. Knöpfel, T., Gallero-Salas, Y. & Song, C. Genetically encoded voltage indicators for large scale cortical imaging come of age. Curr Opin Chem Biol 27, 75–83 (2015).

11. Xiao, S. et al. Large-Scale Voltage Imaging in Behaving Mice Using Targeted Illumination. iScience 103263 (2021) doi:10.1016/j.isci.2021.103263.

12. Kazemipour, A. et al. Kilohertz frame-rate two-photon tomography. Nat Methods 16, 778–786 (2019).

13. Wū, J. et al. Kilohertz two-photon fluorescence microscopy imaging of neural activity in vivo. Nat Methods 17, 287–290 (2020).

14. Abdelfattah, A. S. et al. Sensitivity optimization of a rhodopsin-based fluorescent voltage indicator. 2021.11.09.467909 https://www.biorxiv.org/content/10.1101/2021.11.09.467909v1 (2021) doi:10.1101/2021.11.09.467909.

15. Badon, A. et al. Video-rate large-scale imaging with Multi-Z confocal microscopy. Optica 6, 389–395 (2019).

16. Tsang, J.-M. et al. Fast, multiplane line-scan confocal microscopy using axially distributed slits. Biomed Opt Express 12, 1339–1350 (2021).

17. Packer, A. M., Russell, L. E., Dalgleish, H. W. P. & Häusser, M. Simultaneous all-optical manipulation and recording of neural circuit activity with cellular resolution in vivo. Nat Methods 12, 140–146 (2015).

18. Zhang, Z., Russell, L. E., Packer, A. M., Gauld, O. M. & Häusser, M. Closed-loop all-optical interrogation of neural circuits in vivo. Nat. Methods 15, 1037–1040 (2018).

## METHODS REFERENCES

1. Kim, J.-Y., Grunke, S. D., Levites, Y., Golde, T. E. & Jankowsky, J. L. Intracerebroventricular viral injection of the neonatal mouse brain for persistent and widespread neuronal transduction. J Vis Exp 51863 (2014) doi:10.3791/51863.

2. Ting, J. T., Daigle, T. L., Chen, Q. & Feng, G. Acute Brain Slice Methods for Adult and Aging Animals: Application of Targeted Patch Clamp Analysis and Optogenetics. in Patch-Clamp Methods and Protocols (eds. Martina, M. & Taverna, S.) 221–242 (Springer New York, 2014). doi: 10.1007/978-1-4939-1096-0_14.

3. Abdelfattah, A. S. et al. Bright and photostable chemigenetic indicators for extended in vivo voltage imaging. Science 365, 699–704 (2019).

4. Grimm, J. B. et al. A general method to fine-tune fluorophores for live-cell and in vivo imaging. Nat Methods 14, 987–994 (2017).

5. Chen, T.-W. et al. Ultrasensitive fluorescent proteins for imaging neuronal activity. Nature 499, 295–300 (2013).

6. Peron, S. P., Freeman, J., Iyer, V., Guo, C. & Svoboda, K. A Cellular Resolution Map of Barrel Cortex Activity during Tactile Behavior. Neuron 86, 783–799 (2015).

## REFERENCES

1. Lakowicz, J. R. Principles of fluorescence spectroscopy. (Springer, 2006).

